# Mosquito Olfactory Response Ensemble: a curated database of behavioral and electrophysiological responses enables pattern discovery

**DOI:** 10.1101/2022.01.04.474842

**Authors:** Abhishek Gupta, Swikriti S. Singh, Aarush M. Mittal, Pranjul Singh, Shefali Goyal, Karthikeyan R. Kannan, Arjit K. Gupta, Nitin Gupta

## Abstract

Many experimental studies have examined behavioral and electrophysiological responses of mosquitoes to odors. However, the differences across studies in data collection, processing, and reporting make it difficult to perform large-scale analyses combining data from multiple studies. Here we extract and standardize data for 12 mosquito species, along with *Drosophila melanogaster* for comparison, from over 170 studies and curate the Mosquito Olfactory Response Ensemble (MORE), publicly available at https://neuralsystems.github.io/MORE. We demonstrate the ability of MORE in generating biological insights by finding patterns across studies. Our analyses reveal that ORs are tuned to specific ranges of several physicochemical properties of odorants; the empty-neuron recording technique for measuring OR responses is more sensitive than the *Xenopus* oocyte technique; there are systematic differences in the behavioral preferences reported by different types of assays; and odorants tend to become less attractive or more aversive at higher concentrations.

## Introduction

Mosquitoes find hosts for blood-feeding using various cues, including odors released by the hosts (Degennaro et al., 2013; McBride, 2016; Vinauger et al., 2019). Odorants are detected by sensory neurons located on the peripheral sensory organs, primarily the antennae and the maxillary palps. These neurons express various receptors for detecting odors, including odorant receptors (ORs), gustatory receptors (GRs), and ionotropic receptors (IRs). Sensory neurons transmit information to the antennal lobe in the brain for further processing (Anton et al., 2003; Su et al., 2009).

Because of their relevance to diseases, the olfactory behaviors of mosquitoes have been studied for a very long time (Kline et al., 1990; Mehr et al., 1990; Price et al., 1979). Researchers have employed several types of behavioral assays, such as Y-tube olfactometers, dual-port assays, arm-in-cage landing assays, wind-tunnels, tip assays, and T-mazes (Afify and Potter, 2020; Geier and Boeckh, 1999; Knaden et al., 2012; Logan et al., 2010; Macwilliam et al., 2018; Pates et al., 2001; Simonnet et al., 2014; Spitzen et al., 2013) to quantify the behavior. In parallel, various electrophysiology techniques such as electroantennography, single-sensillum recordings (both in wild-type animals as well as in heterologous expression systems), and voltage-clamp recordings of receptors have been used to quantify the sensory responses to many different odorants (Carey et al., 2010; de Bruyne et al., 2001; De Fouchier et al., 2017; Hallem and Carlson, 2006; Wang et al., 2010). Recent advancements in the techniques to produce transgenic insects have further boosted the research on the mosquito olfactory system (Afify et al., 2019; Kistler et al., 2015; Raji et al., 2019; Riabinina et al., 2016).

The behavioral and electrophysiological data produced from these studies are currently scattered across hundreds of research articles in an unformatted way. Having all this data in one place, in a structured format, can enable systematic large-scale analyses to discover trends that cannot be seen with individual studies (Crasto et al., 2002; Liu et al., 2004, 2011; Marenco et al., 2016; Olender et al., 2013). The Database of Odorant Responses (DoOR) catalogs the OR responses of different odors in *Drosophila melanogaster* (Galizia et al., 2010; Münch and Galizia, 2016) and has proved to be very useful in enabling large-scale computational analyses (Chepurwar et al., 2019; Dasgupta et al., 2018; Saberi and Seyed-Allaei, 2016; Zwicker et al., 2016). However, no such curated dataset is available for mosquitoes. Further, while DoOR only has OR response data, a curated dataset that puts together different kinds of behavioral and electrophysiological recordings would be more powerful.

Here we have compiled the available behavioral and electrophysiological olfactory responses in many species of mosquitoes from over 170 research papers. This curated dataset brings results from diverse sources into a standard format. We have annotated each data-point with various experimental parameters, such as the concentration of the odorant used or the age and the sex of the animals on which the experiments were performed. We demonstrate how the dataset can be used to gain insights into the olfactory system.

## Results

### Curating a comprehensive dataset of olfactory responses

We manually collected a large number of research articles that have reported different kinds of olfactory responses in mosquitoes. The responses were sorted into four different data-types: (1) OR: electrophysiological measurements from genetically labeled odorant receptors, using the empty-neuron system (Carey et al., 2010) or other heterologous expression systems; (2) SSR: single-sensillum recordings without genetic identification of the odorant receptors; (3) EAG: electroantennogram recordings; (4) Behavior: measurements of behavioral preferences to odors.

In most of the articles, the data were reported in text or plots, rather than spreadsheets, and thus had to be manually extracted (see Methods). Preprocessing was often required to convert the data into standard formats: for example, odor preference results could be reported as preference index, percent repellency, percent attraction, etc.; we converted all of them to a common metric – the preference index, calculated as the number of animals choosing the test odor minus the number of animals choosing the control divided by the sum of the two numbers. Similarly, EAG and OR response datasets were processed wherever required to ensure uniformity in data normalization and background subtraction (see Methods).

In total, we collected 30,741 data-points (Fig. 1A), where each data-point corresponds to one of the 4 types of responses for an odorant, covering a total of 758 different odorants (Supplementary Fig. S1A). Care was taken to map the entries reported for different synonyms of the same odorant to a standard name, and to convert the odorant concentrations into standard units (see Methods). We were able to collect data from 12 different species of mosquitoes: *Anopheles gambiae, Aedes aegypti, Culex quinquefasciatus, Anopheles stephensi, Culex pipiens, Aedes albopictus, Culex nigripalpus, Culex tarsalis, Anopheles quadrimaculatus, Anopheles quadriannulatus, Anopheles arabiensis, and Anopheles Coluzzii;* data from *Drosophila melanogaster* was also included to help with comparative analyses (Fig. 1B). The data were sourced from 170 different research papers (Fig. 1C), published over a period of more than 4 decades (Supplementary Fig. S1B).

**Fig. 1:**
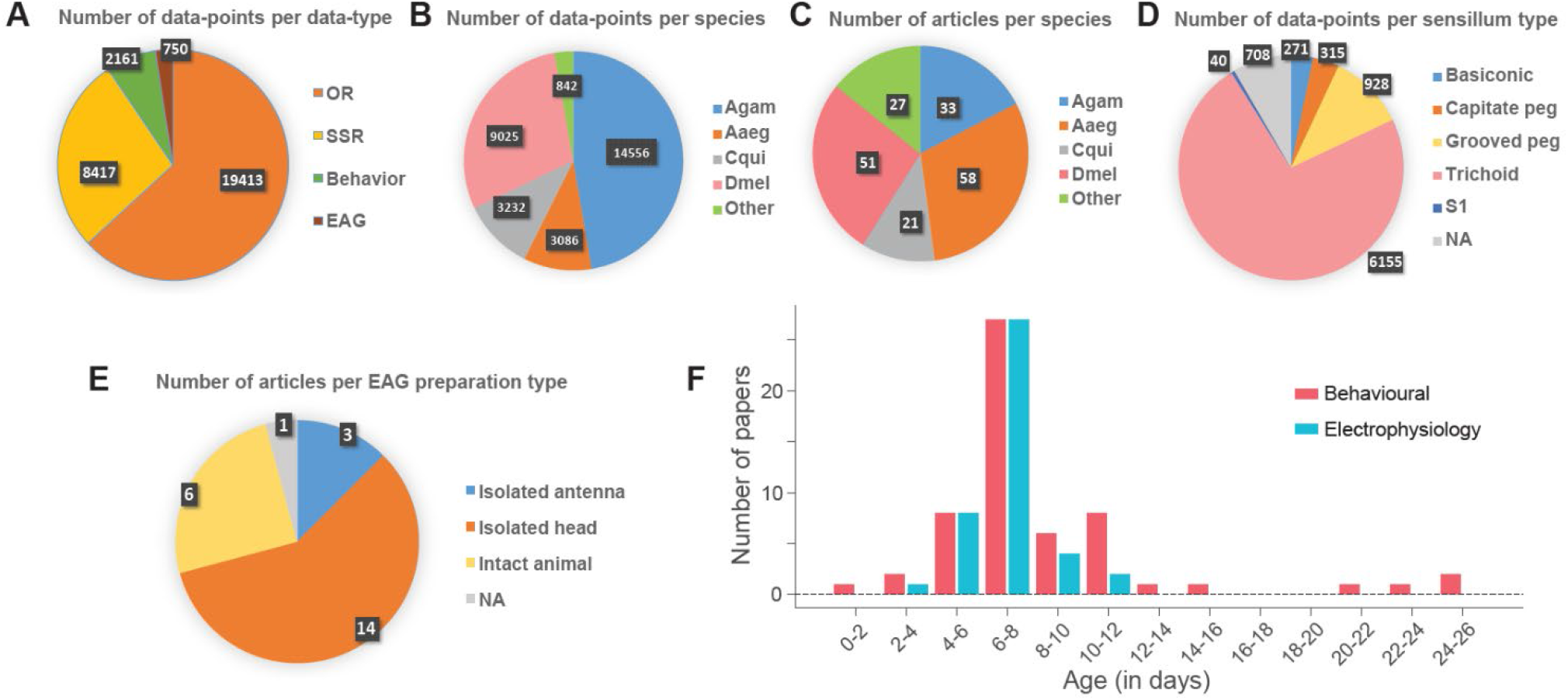
An overview of the dataset. **A** Pie-chart showing the number of data points corresponding to each data type. Each data point corresponds to the response to an odor, and a data type represents the type of response. **B** Pie-chart indicates the number of data points per species. **C** Pie-chart showing the number of studies from which data were extracted for each species. The total number of studies was 170, but some studies included data for multiple species. **D** Pie-chart indicates the number of odor-responses per sensillum type in the Single Sensillum Recording dataset. **E** Pie-chart showing the number of studies that use a particular animal preparation type for Electroantennography recordings. **F** Bars indicate the number of studies verses the age group of the animal for behavioral (red) and electrophysiology (blue) experiments. In **D, E** ‘NA’ category represents not available. Abbreviations: Agam - *Anopheles gambiae*, Aaeg - *Aedes aegypti*, Cqui - *Culex quinquefasciatus*, Dmel - *Drosophila melanogaster*, OR - Odorant Receptor response, SSR – Single Sensillum Recordings, Behavior – Behavioral preference, EAG – Electroantennography

The comprehensive dataset allowed us to note some trends in the experimental preparations. In terms of the number of papers as well as the number of data-points, the three mosquito species with maximum research on the olfactory system are *Anopheles gambiae, Aedes aegypti*, and *Culex quinquefasciatus* (Fig. 1B, C). Among all SSR recordings that are available for mosquitoes, nearly 73% have been performed on the trichoid sensilla (Fig. 1D). Among EAG studies, we noted the experiments have been performed in three kinds of preparations: intact animals, isolated heads, and isolated antenna; the isolated head preparation has been used in more than half of the studies (Fig. 1E). We also checked the ages of mosquitoes used in behavioral and electrophysiological studies and found that most studies have used animals of age 6-8 days for both kinds of experiments, with only a few using younger or older animals (Fig. 1F).

### Web interface for accessing the dataset

We have made the whole dataset available freely through a website: http://neuralsystems.github.io/MORE. The website is organized into 4 sections, corresponding to the 4 data-types. The data are displayed in a tabulated format, which can be sorted in the increasing or the decreasing order of any selected feature (Supplementary Fig. S2). Each row provides one data-point for an odor, along with the corresponding experimental details (such as odor concentration or the species used) and the reference. The row can be expanded to see additional details about the experimental conditions. A search box allows users to enter a term, so that only the rows containing the terms are displayed; this is particularly useful if a user wants to find data for a particular odor, a particular species, or a particular experimental condition, among the thousands of data-points. The results displayed on any screen can be downloaded as an Excel spreadsheet.

The whole dataset can also be downloaded as an Excel file with a single click, without requiring any registration or permissions. This will enable other researchers to use this large and structured dataset, possibly in combination with new or other kinds of data, to conduct new analyses.

### Relationship between OR responses and physicochemical properties of odorant molecules

An analysis of OR responses in *Drosophila* previously suggested that ORs tend to respond more strongly to odorant molecules whose volumes are in a specific range (Saberi and Seyed-Allaei, 2016). Our dataset allowed us an opportunity to systematically examine such relationships between different physicochemical properties and OR responses, and further check if they are conserved between *Drosophila* and mosquitoes. We retrieved the physicochemical properties of odorants from PubChem and analyzed the correspondence between 13 different properties and OR responses (see Methods).

We found that the mosquito ORs responded most strongly to odorants with molecular volumes around 100 Å^3^, in a bell-shaped tuning curve; interestingly, the tuning was largely overlapping between mosquitoes and *Drosophila* (Fig. 2A). To quantify the tuning, we fitted the distribution to a Gaussian and estimated the standard deviation (σ); a smaller value of σ indicates a sharper tuning. To determine the statistical reliability of the observed tuning (with a null hypothesis of no tuning), we compared the observed σ with the values of σ obtained after shuffling the mapping between the responses and the molecular volume (see Methods). This analysis confirmed that the tuning observed for molecular volume was statistically reliable (P < 0.001).

**Fig. 2:**
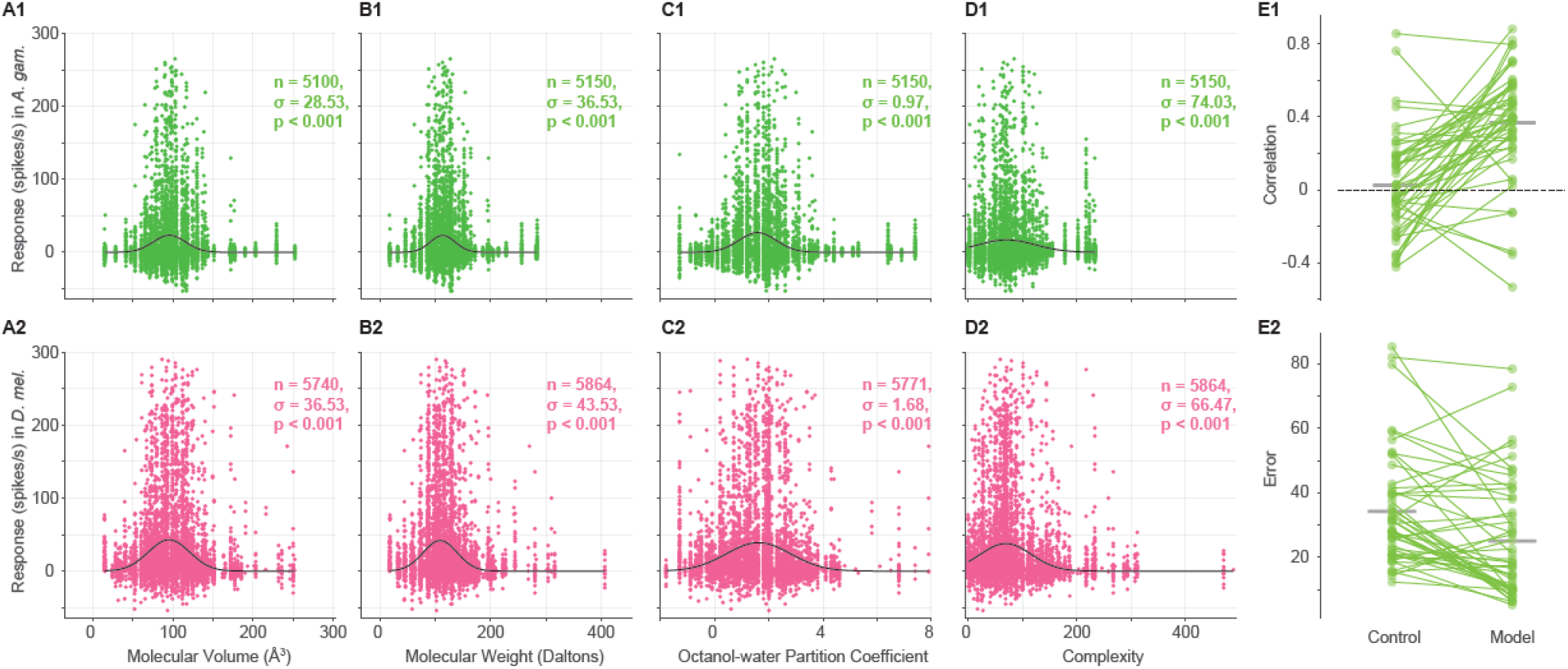
Relationship between OR responses and physicochemical properties of odorant molecules. **A-D** Scatter plots of OR responses (in spikes/second) to odors and the physicochemical properties of those odors, including Volume 3D (**A**), Molecular weight (**B**), Octanol-water partition coefficient (**C**), and Complexity parameter (**D**), in *Anopheles* (**A1, B1, C1, D1**) and *Drosophila* (**A2, B2, C2, D2**). Each point corresponds to an OR-odor pair. In the plots, n is the number of OR-odor pairs, s is the standard deviation of the fitted gaussian, and p is the p-value. In all plots, the black curve corresponds to the fitted gaussian line. The value of n in different plots could differ as the values of all physio-chemical properties were not available for some of the odors. **E** Comparison of the performances of the neural network model and the control model on the test dataset. **E1** shows the correlations between the prediction and the actual odorant responses for each OR; **E2** shows the error (absolute difference between actual and predicted responses) averaged over all test odors. In both plots, each point corresponds to an OR (N = 50).

Similar tuning curves were also observed for molecular weight (Fig. 2B), octanol-water partition coefficient (Fig. 2C), and molecular complexity (Fig. 2D) in *A. gambiae* and *D. melanogaster*. In total, out of the 13 properties examined, we found statically significant response tuning for 12 properties (Supplementary Fig. S3); the only exception was “Conformer Count 3D” (the number of different conformers of the molecule; Supplementary Fig. S3A). Overall, this analysis suggests that insect olfactory systems have evolved to respond preferentially to molecules whose various physicochemical properties lie in certain ranges.

Next, we checked whether a model could be trained to predict the OR responses in *A. gambiae* using the physiochemical properties of odorants. For this analysis, we used a larger set of 295 properties (see Methods). We trained a feedforward neural network model for each OR using 70% of the odors for training, 15% for model validation and keeping 15% for test (see Methods). We found that the responses predicted by the model for the test odors showed higher correlations (R = 0.37 ± 0.30, mean ± s.d.; N = 50 ORs) with the actual responses for the same odors with in an OR, compared the predictions of a control model (R = 0.02 ± 0.28; N = 50 ORs; see Methods); the difference was statistically significant (*P* = 8.73× 10^−7^, sign-rank test; N = 50 ORs; Fig. 2E1). We also checked the magnitude of the errors in the predictions, quantified as the average of the absolute differences between the predicted and the actual responses for the test odors (Fig. 2E2): the error for the model predictions (24.99 ± 17.96, N = 50 ORs) was smaller than the error for the control predictions (34.26 ± 17.24, N = 50 ORs) by 9.27 spikes/s (*P* =1.98× 10^−4^; *N* = 50 *ORs*). These results suggest that machine learning-based models can be used with physicochemical properties of odorants to predict OR responses to novel odorants.

### Differences in techniques for measuring OR responses

Next, we compared different methods for calculating OR responses to odors. In *A. gambiae*, responses of many ORs to a large panel of odors have been measured using two different methods: (1) the empty-neuron system (Carey et al., 2010) in which the OR of interest is expressed in an accessible sensillum of *Drosophila* in place of the native OR, and its response is then measured in the units of spikes/s using the SSR technique; (2) the *Xenopus* oocyte expression system (Wang et al., 2010), in which the OR of interest is expressed in an oocyte, and the response is then measured in the units of nano-Amperes using two-electrode voltage clamp.

By comparing the responses in these two datasets for the same OR-odor combinations, we found that a large fraction (1738 out of 2423; 71.7%) of combinations show non-zero responses in the empty-neuron system and zero responses in the oocyte-recording technique. However, very few combinations (13 out of 2423; 0.5%) show the reverse trend of zero responses in empty-neuron and non-zero responses in oocyte-recording (Fig. 3). To understand the reason for this surprising abundance of zero values in the oocyte recordings, we checked if these mainly correspond to OR-odor combinations that generate an inhibitory (negative) response in the empty-neuron recordings. We found that the zero responses in oocyte-recordings are not limited to cases where the empty-neuron response is negative: in fact, out of 1889 cases with zero responses in oocyte recordings, 918 (48.6%) have a positive response in the empty-neuron recordings, 151 (8%) have zero response, and only 820 (43.4%) have a negative response. Moreover, among the combinations that have negative empty-neuron responses, the responses of these latter 820 combinations with zero oocyte responses are no more negative (−8.55 ± 8.64 spikes/s, mean ± s.d.) than the responses of the 99 combinations with non-zero oocyte responses (−11.32 ± 10.56). Thus, the zero responses in oocyte recordings do not necessarily correspond to inhibitory responses; rather, our analysis of these two datasets suggests that the oocyte recording technique is less sensitive than the empty-neuron technique at detecting OR responses for the same set of odors.

**Fig. 3:**
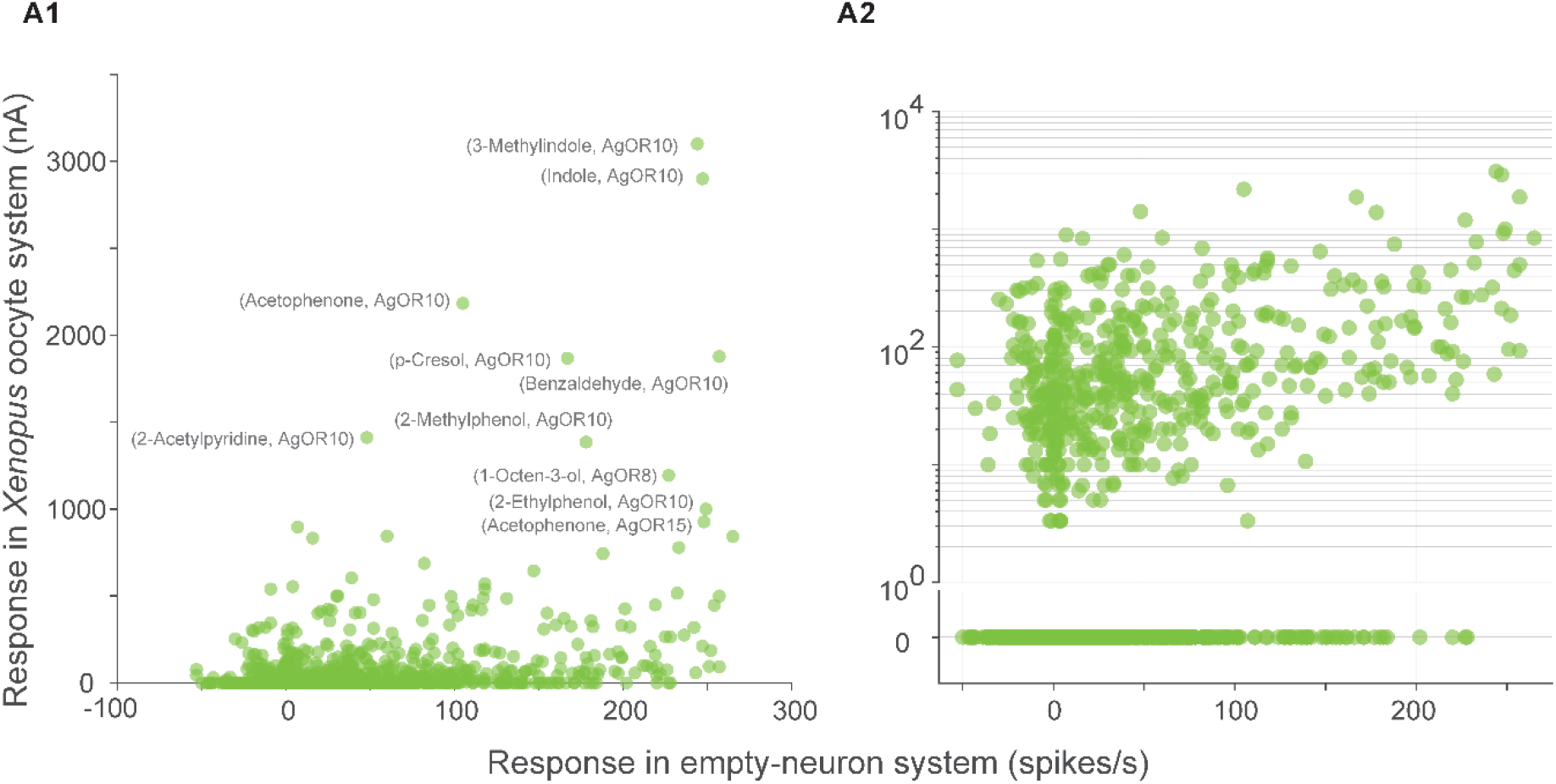
Differences in techniques for measuring OR responses. Scatter plots of OR responses in *Xenopus* oocyte expression system (in units of nano ampere) and empty-neuron system (in units of spikes/second) in linear (**A1**) and logarithmic (**A2**) y-scale (n = 2423 OR-odor measurements). In panel A1, responses greater than 900 in the oocyte system are labeled with the name of the odorant and the OR.

### Differences in behavioral assays

Many different assays have been devised and used by different laboratories to measure the behavioral attractiveness or aversiveness of an odor. Our data collection from multiple studies offers an opportunity to see the most frequently used assays and their relative abundance.

In mosquitoes, these assays belonged to five broad categories: Y-tube (Geier et al., 1996), dual-port (Pates et al., 2001), wind-tunnel (Healy and Copland, 2000), tip (Afify and Potter, 2020), and landing (or arm-in-cage) assays (Ali et al., 2017; Logan et al., 2010) (Fig. 4A1). Although all these assays quantify the preference of mosquitoes to the tested odor, they can differ in the odorant exposure profile and the specific motor actions used by the mosquitoes: for example, a Y-tube assay involves a choice between two alternatives in a confined chamber, a wind-tunnel involves free flight movement towards an odor source in a large chamber, and a landing assay involves the termination of flight followed by landing very close to an odor source. For adult mosquitoes, Y-tube assay was the most abundant (33.3%), followed by dual-port (28.6%) and landing assays (28.6%), and wind-tunnel (7.9%) and tip assay (1.6%) were the least abundant. In *Drosophila*, the assays could be grouped into three categories: Y-maze (Charro and Alcorta, 1994), T-maze (Helfand and Carlson, 1989), and dual-port (or trap) assays (Knaden et al., 2012) (Fig. 4A2). Among these, T-maze assays were the most common (54.6%) in our dataset, followed by Y-maze (24.2%) and dual-port (21.2%) assays.

**Fig. 4:**
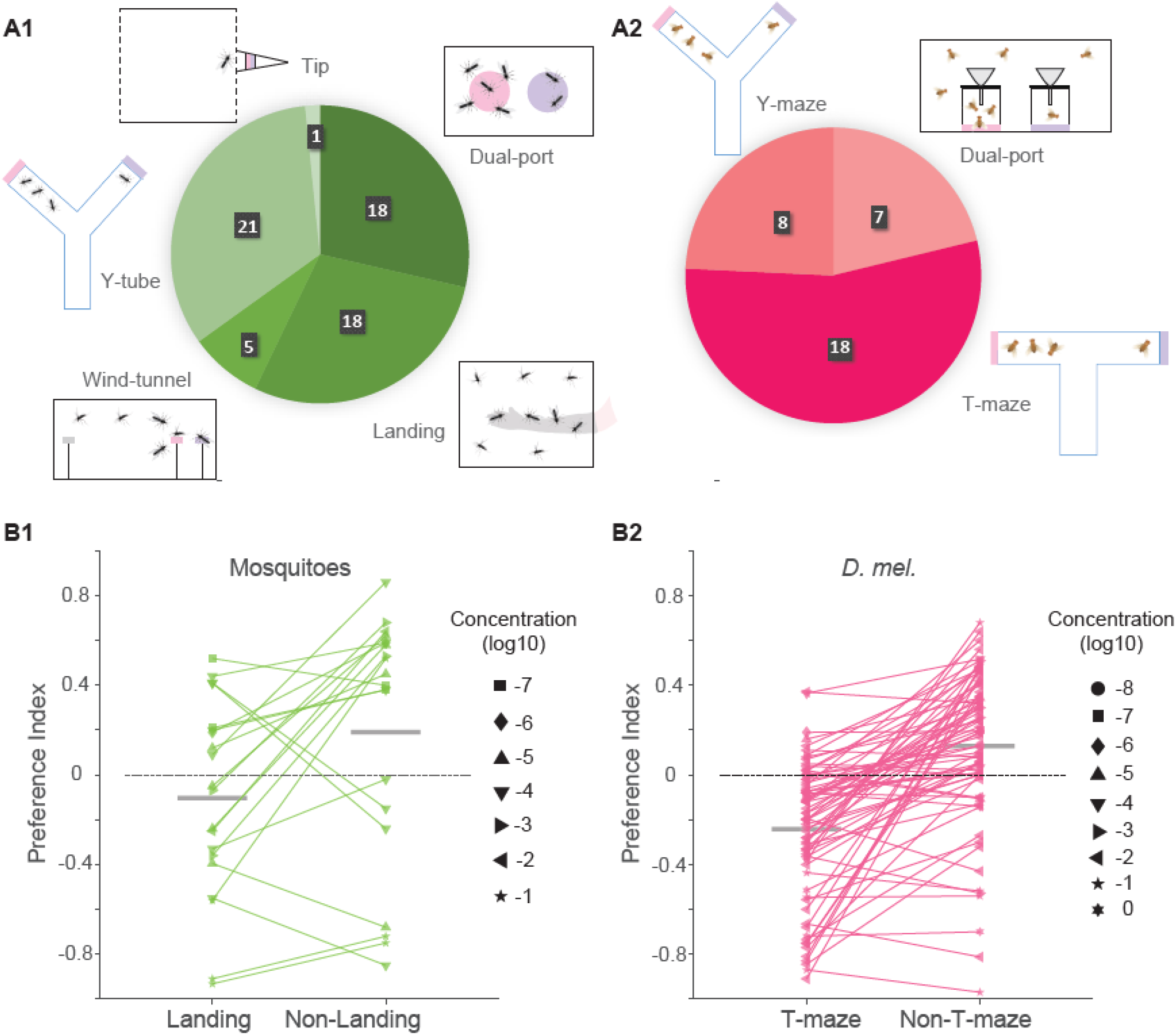
Differences in behavioral assays. **A1, A2**, Pie-charts showing the number of studies from which data were extracted for different behavioral assays in mosquitoes (**A1**) and *Drosophila* (**A2**). **B1, B2**, Comparison of the preference indices in landing and non-landing (Dual-port and Y-tube) assays in mosquitoes (**B1**), and T-maze and non-T-maze (Dual-port and Y-maze) assays in *Drosophila* (**B2**). Each point represents an odor-concentration pair (n = 20 for mosquitoes and n = 74 for *Drosophila*). The size of each dots indicates the odor concentration (see legend). Horizontal gray lines represent means.

The behavioral attraction or aversion for an odor, estimated from these assays, is often reported as a preference index. Although some variability is expected in the preference indices measured in different studies, because of experimental noise or minor differences in the experimental conditions, it is not known if there are systematic biases in the preference indices reported by different types of assays. We used our large dataset to explore the possibility of such systematic differences. In mosquitoes, we noticed that the preference indices obtained in the landing assays were often smaller (or more negative) than the indices for the same odors in other types of assays (*P* = 1.68 × 10^−2^, *N* = 20 pairs of data points from *Aedes aegypti* and *Culex quinquefasciatus*; Fig. 4B1). In *Drosophila*, we found that T-maze assays reported odors to be more aversive than Y-maze or dual-port assays (*P* = 6.03 × 10^−11^, *N* = 74 pairs of odors; signrank test; Fig. 4B2). These results highlight the need for caution when comparing behavioral preferences of odors across different studies.

### Relationship between the preference index and the oviposition index

Behavioral preferences are governed by the internal states of the animals (Sayin et al., 2018). There are examples where the attraction or aversion to an odor during foraging behavior is different from that during egg-laying behavior. For example, *D. melanogaster* show avoidance to acetic acid in odor choice assays during foraging, but attraction to acetic acid during egg-laying (Joseph et al., 2009). Another study found that valencene, b-caryophyllene, b-caryophyllene oxide, and limonene oxide had very different preference indices (during foraging) and oviposition indices in *D. melanogaster* (Dweck et al., 2013).

Our large collection of behavioral data allowed us to systematically examine whether the odor preferences during oviposition are independent of odor preferences during foraging or host-seeking. We selected odors for which both the oviposition index and the preference index were available in the dataset (see Methods), and then compared the two values (Fig. 5). We found that the oviposition index was not correlated with the preference index during host-seeking in *A. aegypti* (*R* = 0.06, *P* = 0.86, *N* = 11; Fig. 5A1) and the preference index during foraging in *D. melanogaster* (*R* = 0.06, *P* = 0.78, *N* = 22; Fig. 5A2). Thus, the preferences during foraging or host-seeking appear completely independent of the preferences during oviposition.

**Fig. 5:**
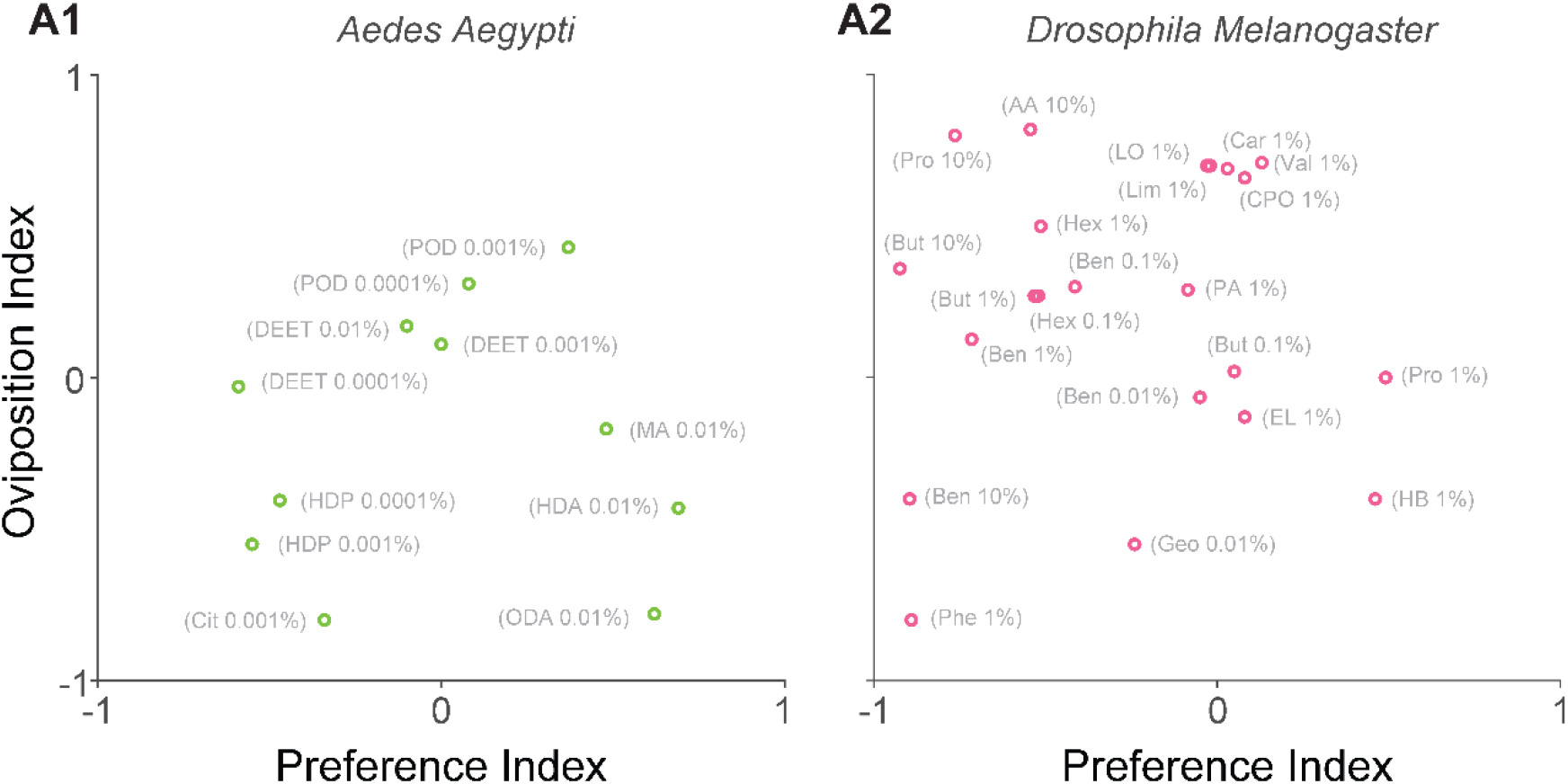
Relationship between the preference index and the oviposition index. **A1, A2**, Scatter plots between the preference index and the oviposition index in *Aedes Aegypti* (**A1**) and *Drosophila* (**A2**). Each point represents an odorant at a specific concentration (n = 11 for mosquitoes and n = 22 for *Drosophila*). Odorant acronyms: DEET, N,N-diethyl-meta-toluamide; POD, Propyl Octadecanoate; MA, Myristic Acid; HDP, Hexadecyl Pentanoate; Cit, Citronellal; HDA, Hexadecanoic Acid; ODA, Octadecanoic Acid; Pro, 1-Propanol; AA, Acetic Acid; But, 1-Butanol; EL, Ethyl Lactate; Ben, Benzaldehyde; Hex, (E)-2-Hexenal; Phe, Phenol; Geo, Geosmin; HB, Hexylbutyrate; PA, Pentanoic Acid; CPO, (−)-Caryophyllene Oxide; LO, Limonene Oxide; Lim, Limonene; Val, Valencene; Car, B-Caryophyllene;

### Comparison of behavioral preference across mosquito species

Our dataset including multiple species of mosquitoes provided an opportunity to check how similar or different are the preferences indices of odors between the species. For each pair of species, we selected odors that were tested in both the species at similar concentrations (see Methods), and calculated the correlation coefficient between their preference indices in the two species. We found weak to moderate correlations in different pairs of species, with the Pearson correlation coefficient varying between 0.12 to 0.82 (Supplementary Fig. S4). The correlation between *A. gambiae* and *A. aegypti* was 0.68 (n =44), while it was 0.59 between *C. quinquefasciatus* and *A. aegypti* (n = 117) and 0.46 between *A. aegypti* and *A. albopictus* (n = 24). We note that the correlation values are affected by the exact identities of the available common odors, which differed for different pairs.

### Dependence of behavioral preference on odor concentration

The concentration of an odor can affect the behavioral preference. In *D. melanogaster*, there are examples where a 10-fold change in concentration can result in either an increase or a decrease in the preference index. Figure 6A1 shows the preference indices of benzaldehyde with concentrations varying over 5 orders of magnitude in T-maze assays: higher concentrations typically show more aversion. Figure 6A2 shows the preference indices of ethanol with concentrations varying over 4 orders of magnitude in Y-maze assays: here, the preference increases from 10^−3^ to 10^−1^, but decreases if the concentration is further increased.

**Fig. 6:**
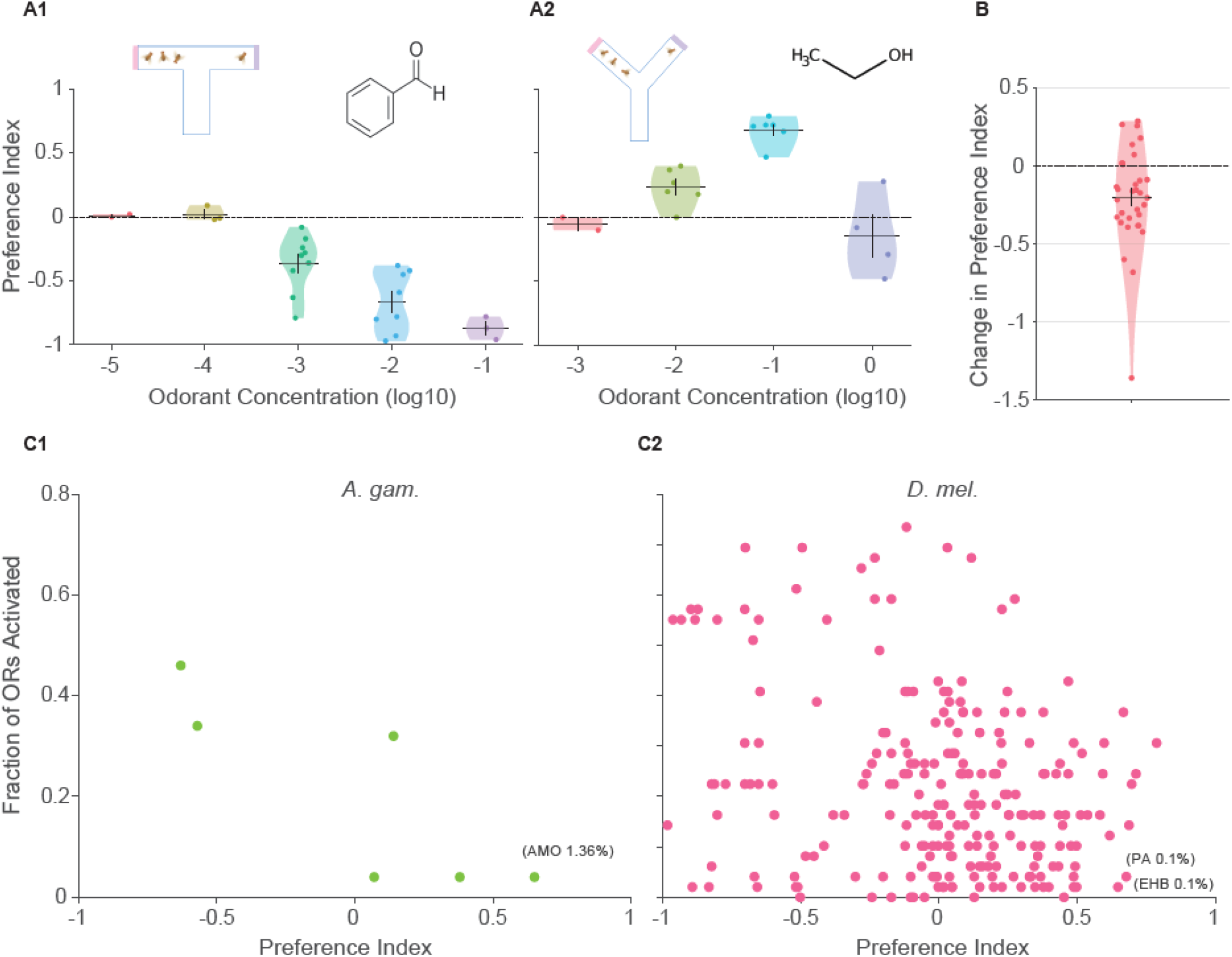
Dependence of behavioral preference on odor concentration. **A1, A2**, Violin plots show the preference indices for different concentrations of benzaldehyde (**A1**) and ethanol (**A2**) in *Drosophila*. Each point within a violin represents the preference index obtained from a different study. **B**, The plot shows the change (mostly reduction) in the preference index on increasing the concentration by ten folds. Each point represents the preference index at a particular concentration minus the preference index at a ten-fold lower concentration of the same odor, in the same type of assay and same species. The plot includes data for n = 33 concentration pairs from *A. aegypti, A. albopictus, A. gambiae, C. pipiens*, and *D. melanogaster*. Odorant acronyms: AMO, Ammonia; PA, Propanoic Acid; EHB, Ethyl-3-hydroxybutyrate. In all plots, the horizontal line and the error bar indicate the mean and the SEM, respectively.

To check if there may be a general pattern in how the preference index varies in response to increasing odor concentration, we collected pairs of preference indices at concentrations separated by a factor of 10 for the same odor, using the same type of assay in the same species (Supplementary Fig. S6; see Methods). In 33 such pairs available in our dataset, we checked the difference between preference index at the higher concentration and at the lower concentration (Fig. 6B). We found that increasing the odor concentration 10-folds decreases the preference index, on average by 0.2 (*P* = 4.01 × 10^−4^, *N* = 33; *signed rank test*). The same trend was observed even when we limited the analysis to only those pairs where the preference index at the lower concentration was negative (*mean change* = −0.14, *P* = 0.04, *N* = 11; Supplementary Fig. S5A1) or positive (*mean change* = −0.23, *P* = 0.005, *N* = 22; Supplementary Fig. S5A2). Thus, our analysis using this large dataset suggests that an increase in odor concentration tends to make aversive odors more aversive and attractive odors less attractive.

To probe this further, we checked the relationship between the number of ORs activated (increase of at least 10 spikes) by an odor and the preference index of the odor. This analysis revealed a negative correlation between the two parameters in both *A. gambiae* (*R* = −0.83, *P* = 0.039, *N* = 6; Fig. 6C1) and *D. melanogaster* (*R* = −0.27, *P* = 3.7 × 10^−5^, *N* = 230; Fig. 6C2). As higher concentrations are likely to activate non-specific ORs, this provides a possible explanation for why higher concentrations tend to be more aversive. We further highlight the odors that activated a small fraction of the ORs and were highly attractive. In *A. gambiae*, ammonia 1.36% activates AgOR46 and AgOR50 and has a preference index of 0.65. In *D. melanogaster*, propanoic acid 0.1% activates OR24a and OR42a and has a preference index of 0.68, and ethyl-3-hydroxybutyrate 0.1% activates OR85a and has a preference index of 0.65.

## Discussion

In summary, we have curated MORE, a large dataset of behavioral and electrophysiological responses in 12 species of mosquitoes along with *D. melanogaster*. The dataset includes 30,741 data-points for 758 odorants collected from 170 research articles.

Bringing this scattered information into a well-structured database involved several challenges, because of the different preprocessing steps or the different units or metrics used by the studies. Some studies normalized the EAG responses using the responses to a reference odor, while others reported the raw values. Some studies reported the OR electrophysiological responses after subtracting the background activity, while others reported without this step. In MORE, we processed the data to include uniform normalization and background subtraction. Odor preferences in behavioral studies were reported using a variety of metrics, such as preference index (Yu et al., 2015), percent attraction (Geier et al., 1996), percent repellency (Islam et al., 2017b), protective efficacy (Logan et al., 2010), etc. In MORE, we converted the reported preferences in all papers into the common metric of preference index. The odor concentrations were reported in a variety of units, which had to be standardized before their inclusion in MORE. Different studies referred to the same odorants using different names. For example, isoamyl alcohol, isopentyl alcohol, and isopentanol, all are common names of 3-methyl-1-butanol. In MORE, we combined all such data-points using a single standard name for each odorant. In many studies, the data-points were not available in an accessible format, and had to be obtained either by requesting the original authors or by extracting from the figures using a computer script. We have focused on mono-molecular odorants currently; future work may explore how to incorporate odor blend (Mozūraitis et al., 2020).

The structured format of MORE makes the data amenable to large-scale analyses of patterns across the datasets, as we have demonstrated here. MORE can also facilitate the application of machine learning methods that are particularly dependent on large and structured datasets. We have created an interactive website for browsing the data, while also providing an easy option for downloading the entire dataset for offline analyses.

We found that in mosquitoes as well as in flies, the sensory responses were tuned to specific ranges of various physicochemical properties of the odorant molecules. The knowledge of these ranges may be useful in designing synthetic agonists for the ORs. We observed reliable tuning for 12 of the 13 physicochemical properties we tested. One of these properties, the octanol-water partition coefficient, is known to be related to the air-mucus odorant partition coefficient (Scott et al., 2014). Another property, molecular complexity, has been reported to be the determinant of the number of olfactory notes and the pleasantness of smell (Kermen et al., 2011). The one property that did not show tuning was the number of different 3D conformers of the molecule – this is not surprising as this particular property informs about the possible variations in the molecule but does not tell about the shape of any specific structure, unlike the other 12 properties. We also observed that the molecular properties of the odorants could be used to train a neural network model for the predicting the OR responses to new odorants. The accuracy of these predictions is likely to improve as more data becomes available for training the model.

We found systematic differences in the OR responses recorded using the empty-neuron system and the *Xenopus* oocyte expression system. Our results generalize the experimental observation made by Wang *et al*. using one pheromone receptor (Or13) in *Helicoverpa assulta* (Wang et al., 2016). The sensitivity of the two techniques might differ due to differences in the levels of receptor expression in the different kinds of cells, and differences between the odor delivery through the liquid medium in the oocyte recording technique and the volatile odor delivery in the empty-neuron technique. These results highlight the need for caution when interpreting negative results from the oocyte expression system.

Our dataset revealed no correlation between the oviposition indices and the preference indices of odors in mosquitoes or *Drosophila*. This result is consistent with previous work showing that sensory processing and the choice of behavior are expected to be state-dependent (Barrozo et al., 2011; Cohn et al., 2015; Gadenne et al., 2016; Sayin et al., 2018; Vogt et al., 2021). We also found that higher odor concentrations were in general more aversive than lower concentrations of the same odorants (Fig. 6). This effect may also be related to our observations that landing assays resulted in lower or more negative preference indices than the non-landing assays for mosquitoes and that T-maze assays resulted in more negative preference indices than non-T-maze assays for flies (Fig. 4): we speculate that these differences could be because the landing assays bring mosquitoes closer to the odor source and expose them to the odorants with less air dilution, and perhaps the size and shape of the T-maze expose flies to higher concentrations than Y-maze assays. In *Drosophila*, a low concentration of apple cider vinegar triggers attraction through a smell set of activated glomeruli, but a higher concentration triggers aversion through activation of an additional glomerulus (Semmelhack and Wang, 2009). Our results show that this concentration-dependent aversion is a more general pattern extending across odors and species.

## Methods

### Data Extraction

Tabulated data from some papers were obtained by requesting the original authors (Hallem and Carlson, 2006; Hill et al., 2009; Wang et al., 2010). From some papers, when the data were not available directly, WebPlotDigitizer tool (provided by Ankit Rohatgi) was used to manually extract the data from plots.

Different papers have reported odorant concentrations using many different formats such as volume/volume (V/V), weight/volume (W/V), molarity, parts per million (ppm), or weight/area (mg/cm^2^), which makes the comparisons difficult. Wherever possible, we converted the concentrations to common notations and units, as either fractions (V/V or W/V) or to g/ml (W/V); in case of dry odorant applied on a filter paper, we mentioned the amount of odorant use after setting concentration type as “Dry”. To take a few examples of the conversions used: 1.11×10^−5^ M of lactic acid was converted to 0.000001 g/ml (Braks et al., 2001); 1ppm was converted to 1mg/l (Islam et al., 2017a); in one study, 0.025 ml of 0.01 mg/cm^2^ odor was added on 6.6cm^2^ cloth, which was converted to equivalent W/V concentration in g/ml given by (0.000001 g/cm^2^) × (6.6 cm^2^) / (0.025 ml) (Mehr et al., 1990).

The EAG responses reported in some studies were not normalized (Cork and Park, 1996; Guha et al., 2014), but in other studies were normalized with respect to a reference odor, such as 1-octen-3-ol (Blackwell and Johnson, 2000; Constantini et al., 2001). To maintain uniformity, if the responses were not normalized, we normalized them with 1-octen-3-ol, if present, or with another suitable reference odor depending on the dataset; the reference odor used is noted for each EAG data-point.

In OR electrophysiological measurements, some studies report the responses after subtracting the spontaneous firing rate (Hallem and Carlson, 2006), while some report the raw response without the background subtraction (Dobritsa et al., 2003). In the latter cases, we subtracted the response to the solvent (e.g., paraffin oil) from the odor responses of each OR.

### Physicochemical properties

The properties of the odorants were obtained from the PubChem database using a MATLAB script (created by Vincent Scalfani). We could obtain a set of 28 properties by this automated approach. Some of the properties were missing for some of the odorants. To ensure that the curve-fitting in the subsequent steps can be done reliably, we analyzed only those properties which took at least ten different numerical values among our set of odorants, leaving us with the following 13 properties: Molecular Weight; Molecular Volume; Octanol-water Partition Coefficient (computationally predicted); Complexity (an indicator of the complexity of the molecular structure, calculated using the Bertz/Hendrickson/Ihlenfeldt formula); Conformer Count 3D (the number of conformers); Effective Rotor Count 3D (number of effective rotors); Feature Count 3D (total number of 3D features); Heavy Atom Count (number of non-hydrogen atoms); Rotatable Bond Count (number of rotatable bonds); Topological Polar Surface Area (estimate of the area that is polar); XSteric Quadrupole 3D (x component of the quadrupole moment); YSteric Quadrupole 3D (y component of the quadrupole moment); ZSteric Quadrupole 3D (z component of the quadrupole moment).

The response data were taken from experiments that used the concentration of 10^−2^ V/V or W/V, which was the most frequently used concentration in the dataset. To calculate the p-values, we first shuffled the mapping between the odorant property and response for 1000 times and then calculated the p-value as the fraction of times σ_shuffle_ ≤ σ_actual_ (where, σ_shuffle_ and σ_actual_ are the standard deviations of fitted Gaussian on the shuffled and actual data, respectively). For fitting the gaussian, we used the MATLAB ‘fit’ function with ‘gauss1’ argument.

### Neural network model for OR response prediction

We obtained 1842 physicochemical descriptors from PubChem and Mordred (Moriwaki et al., 2018). From these descriptors, we first removed the descriptors with missing values or variance <0.005, leaving 645 descriptors. Next, we removed the descriptors with mutual correlation greater than 0.95 (Caballero-Vidal et al., 2020). This finally gave us 295 descriptors, which were used to represent odorants in the model. For predicting the OR responses to the odorants, we used the Deep Learning Toolbox in MATLAB with default parameters to implement a feedforward artificial neural network with three hidden layers, each having 50 neurons. The dataset including 112 odorants for each OR, which were randomly partitioned into a training set (70% of the odorants), a model validation set (15% of the odorants), and a test set (15% of the odorants). The model performance is reported using only the predictions on the test odorants. For comparison, we randomly shuffled the OR-odorant response matrix and used the shuffled responses as the control predictions.

### Analyses of behavioral assays

The exact behavioral assay used in each study is mentioned with the corresponding data-points in the database. In all analyses (Fig. 4, 5, 6), only those data-points were used for which the odor concentration was known. The concentrations were rounded to the nearest power of 10. In the comparison of preference index and oviposition index, the two indices were compared only when both the indices were measured at the same concentration (after rounding). After applying this criterion, we had 11 data-points in *A. aegypti* (shown in Fig. 5); species with less than 5 data-points each were not analyzed. In Figure 4, if multiple data-points from different studies were available for an odor for the same assay category (example: T-maze or non-T-maze) and the same concentration, they were averaged into a single data-point. In Supplementary Figure S4, an odor was included only if it was tested in the compared species at the same concentration (if the same concentration was not available, the closest available concentration within ±10 folds was taken). The correlations were analyzed for only those pairs of species for which at least 10 common odors were available. In Figure 6A-B, we considered an odor-concentration if at least two data-points were available for that concentration with the same assay in the same species (they were averaged into a single data-point).

### Statistical analysis and code availability

All the analyses were conducted in MATLAB. The Gramm plotting toolbox (Morel, 2018) was used to draw the plots. The non-parametric Wilcoxon signed rank test was used to calculate the p-values of paired sample comparisons. The code developed in this study can be accessed from the lab GitHub repository (https://github.com/neuralsystems/MORE).

## Acknowledgments

We would like to thank Elissa A Hallem (University of California, Los Angeles), John Carlson (Yale School of Medicine), Laurence J. Zwiebel (Vanderbilt University), and Sharon Hill (Swedish University of Agricultural Sciences) for sharing raw data from their papers (Hallem and Carlson, 2006; Hill et al., 2009; Wang et al., 2010). We thank members of the N.G. lab for helpful discussions. This work was supported by the DBT/Wellcome Trust India Alliance Fellowship [grant number IA/I/15/2/502091] and the SERB Core Research Grant [CRG/2020/004719] awarded to N.G.

## Author Contributions

AG and NG designed research; AG, SSS, AMM, PS, SG, KRK and AKG collected data; AG performed data analysis in consultation with NG; AG and NG wrote the manuscript with inputs from all authors.

