## Supplementary Figures for "Mosquito Olfactory Response Ensemble: a curated database of behavioral and electrophysiological responses enables pattern discovery"

**Figure S1**

**A**

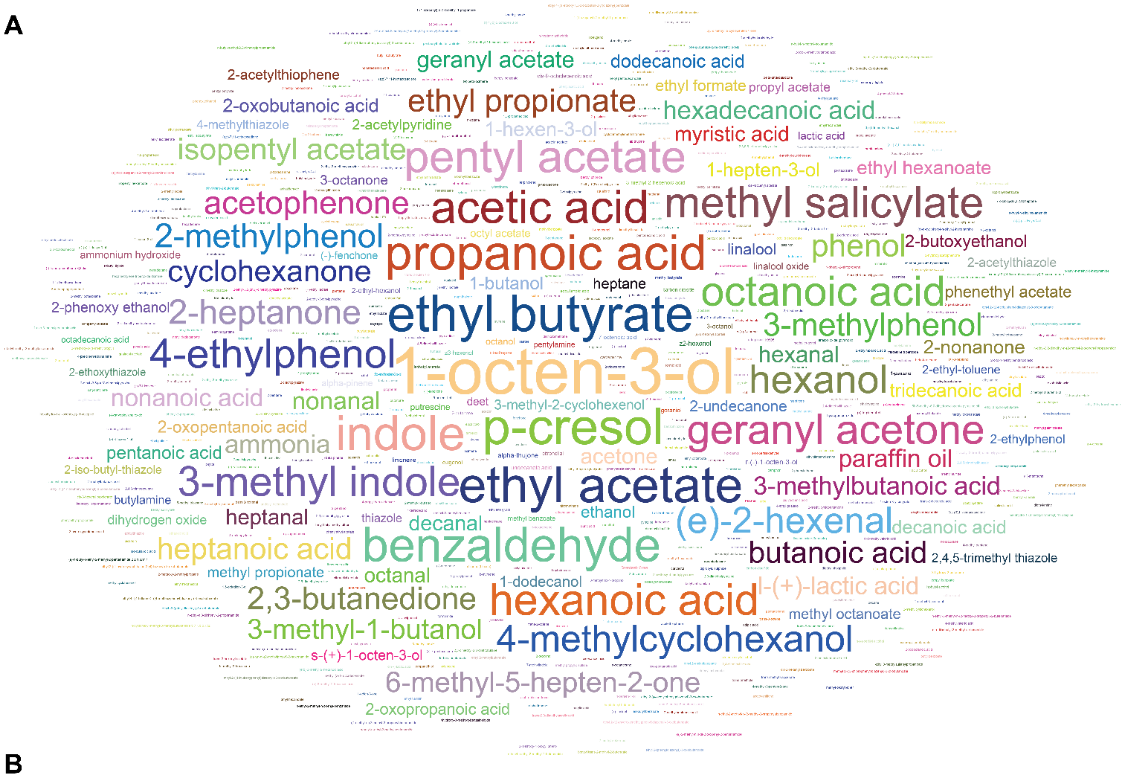


**B**

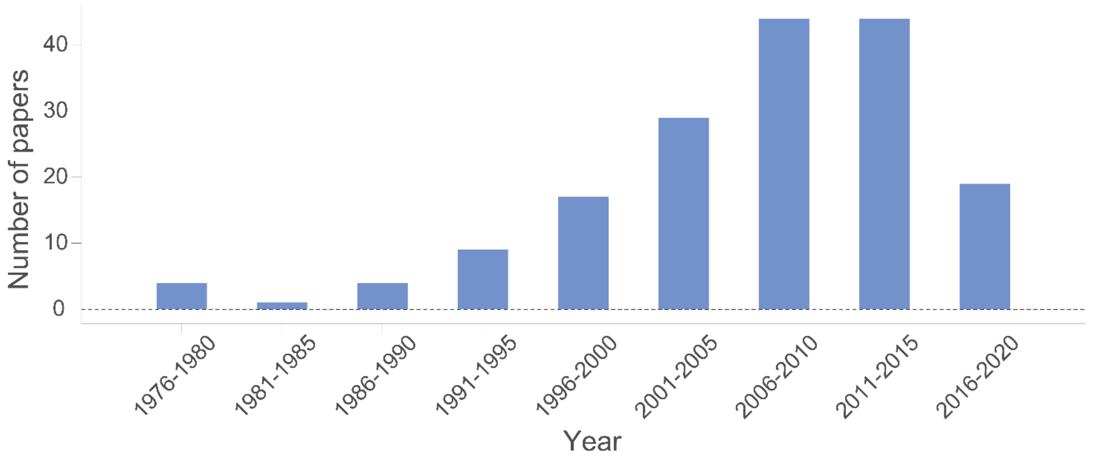


**Fig. S1: A summary of the dataset.**

**A,** Word cloud showing the odor molecules that are present in the dataset. The size of the text is directly proportional to the number of data-points available for that odor in the dataset. In total, the dataset contains responses for 758 odor molecules. The top 10 odorants with the largest number of data-points (in decreasing order) are: 1-Octen-3-ol, p-Cresol, Ethyl butyrate, Propanoic Acid, Indole, Benzaldehyde, Acetic acid, Pentyl acetate, and Hexanoic acid.

**B,** Bars indicate the number of studies included from different years (publication date) in the dataset.

**Figure S2**

Home | Behavior ▾ | Single Sensillum Recording | Electroantennography | Odorant Receptor ▾

Mosquito Preference Index | Drosophila Preference Index | Mosquito Oviposition Index | Drosophila Oviposition Index

Show 10 entries

Search/ Filter

| | odor | concentration | species | assay | response | reference |
|---|---|---|---|---|---|---|
| ⊕ | (+)-alpha-pinene | NA | CQui | Dual-port | 0.27 | Allan2006 |
| ⊕ | (-)-trans-p-Menthane-3,8 diol | NA | AAeg | Landing | R | Ali2017 |
| ⊕ | (E)-1-(1-azepanyl)-2-methyl-2-penten-1-one | NA | AAeg | Landing | R | Katritzky2010 |
| ⊕ | (E)-2-hexenal | 0.000001 | CPip | Y-tube | 0.38 | Yu2015 |
| ⊕ | (E)-2-hexenal | 0.000002 | CPip | Y-tube | 0.12 | Yu2015 |
| ⊕ | (E)-2-hexenal | 0.00000175 | CPip | Y-tube | 0.12 | Yu2015 |
| ⊕ | (E)-2-hexenal | 0.0000015 | CPip | Y-tube | 0.2 | Yu2015 |
| ⊕ | (E)-2-hexenal | 0.00000125 | CPip | Y-tube | 0.12 | Yu2015 |
| ⊕ | (E)-2-hexenal | 0.000001 | CPip | Y-tube | 0.28 | Yu2015 |
| ⊕ | (E)-2-hexenal | 7.5e-7 | CPip | Y-tube | 0.28 | Yu2015 |

Showing 1 to 10 of 1,109 entries | Download filtered data | Previous 1 2 3 4 5 ... 111 Next

**Fig. S2: MORE website**

Screshot shows one of the pages on the MORE website.

**Figure S3**

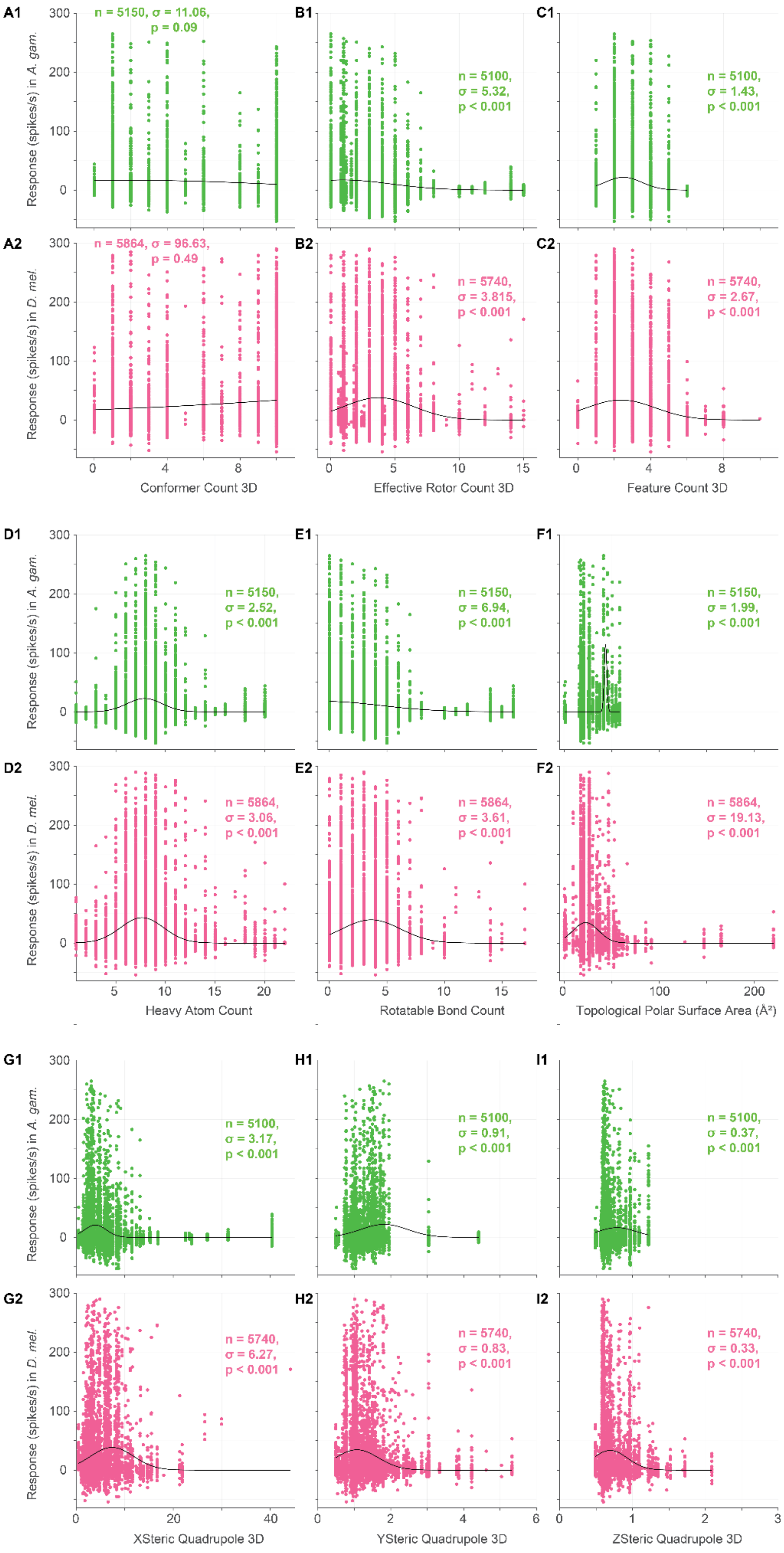

A1
n = 5150, σ = 11.06, p = 0.09
B1
n = 5100, σ = 5.32, p < 0.001
C1
n = 5100, σ = 1.43, p < 0.001
A2
n = 5864, σ = 96.63, p = 0.49
B2
n = 5740, σ = 3.815, p < 0.001
C2
n = 5740, σ = 2.67, p < 0.001
Response (spikes/s) in A. gam.
Response (spikes/s) in D. mel.
Conformer Count 3D
Effective Rotor Count 3D
Feature Count 3D
D1
n = 5150, σ = 2.52, p < 0.001
E1
n = 5150, σ = 6.94, p < 0.001
F1
n = 5150, σ = 1.99, p < 0.001
D2
n = 5864, σ = 3.06, p < 0.001
E2
n = 5864, σ = 3.61, p < 0.001
F2
n = 5864, σ = 19.13, p < 0.001
Heavy Atom Count
Rotatable Bond Count
Topological Polar Surface Area (Å²)
G1
n = 5100, σ = 3.17, p < 0.001
H1
n = 5100, σ = 0.91, p < 0.001
I1
n = 5100, σ = 0.37, p < 0.001
G2
n = 5740, σ = 6.27, p < 0.001
H2
n = 5740, σ = 0.83, p < 0.001
I2
n = 5740, σ = 0.33, p < 0.001
XSteric Quadrupole 3D
YSteric Quadrupole 3D
ZSteric Quadrupole 3D

**Fig. S3: Relationship between OR responses and additional physicochemical properties of odor molecules.**

Scatter plots of OR responses (in spikes/second) to odors and the physicochemical properties of those odors, including Conformer Count 3D (**A**), Effective Rotor Count 3D (**B**), Feature Count 3D (**C**), Heavy Atom Count (**D**), Rotatable Bond Count (**E**), Topological Polar Surface Area (**F**), XSteric Quadrupole 3D (**G**), YSteric Quadrupole 3D (**H**), ZSteric Quadrupole 3D (**I**) in mosquitoes (**A1**, **B1**, **C1**, **D1, E1, F1, G1, H1, I1**) and *Drosophila* (**A2**, **B2**, **C2**, **D2, E2, F2, G2, H2, I2**). Each point corresponds to an OR-odor pair. In the plots, n is the number of OR-odor pairs, σ is the standard deviation of fitted gaussian, and p represents the p-value.

In all the plots, the black line corresponds to the fitted gaussian.

Figure S4

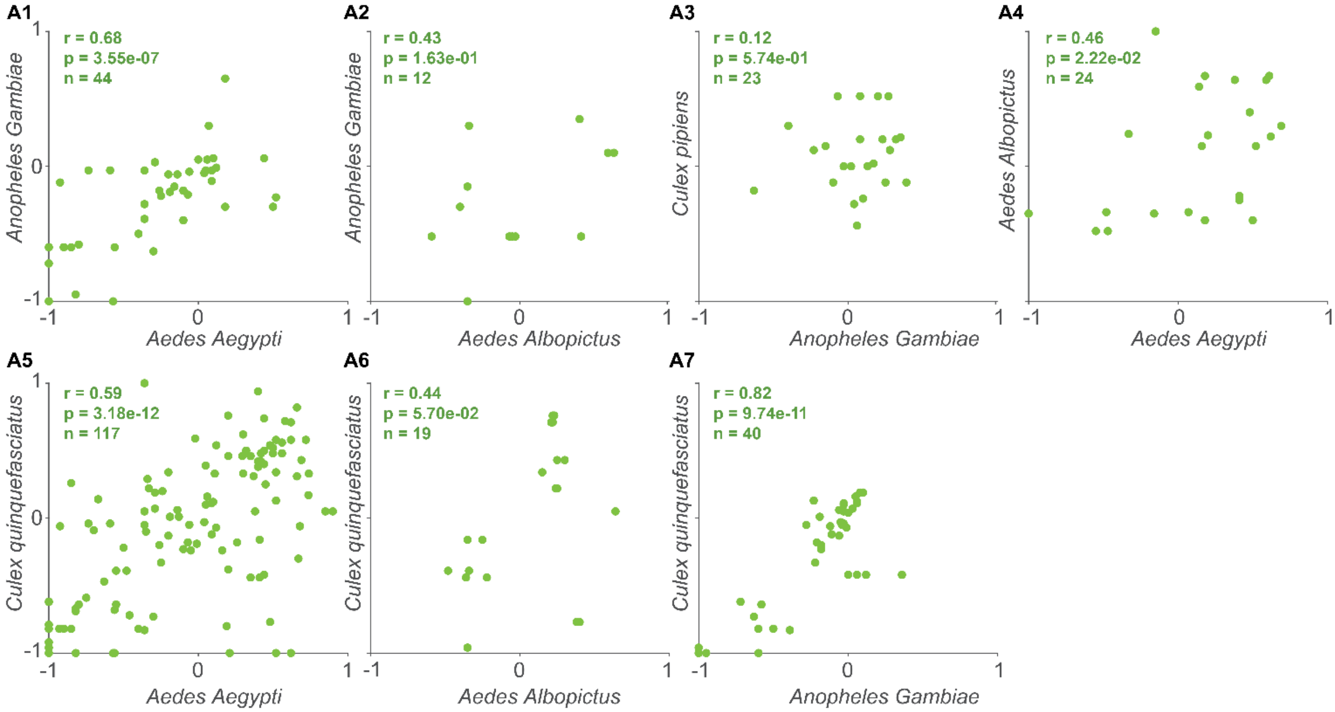


**Fig. S4: Behavioral preferences across different mosquito species**

Scatter plots of preference indices in pairs of mosquito species.

Figure S5

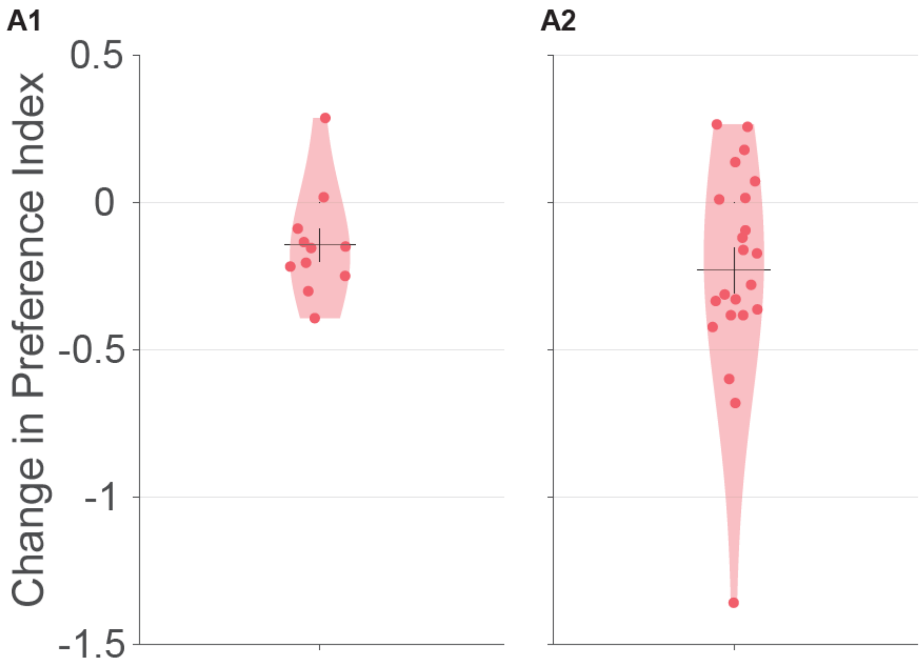


**Fig. S5: Dependence of behavioral preference on odor concentration, for positive or negative preference indices at the lower concentration.**

**A1, A2,** The change in preference index on increasing the concentration by ten folds when the preference index at the lower concentration is negative (**A1**) or positive (**A2**).

Figure S6

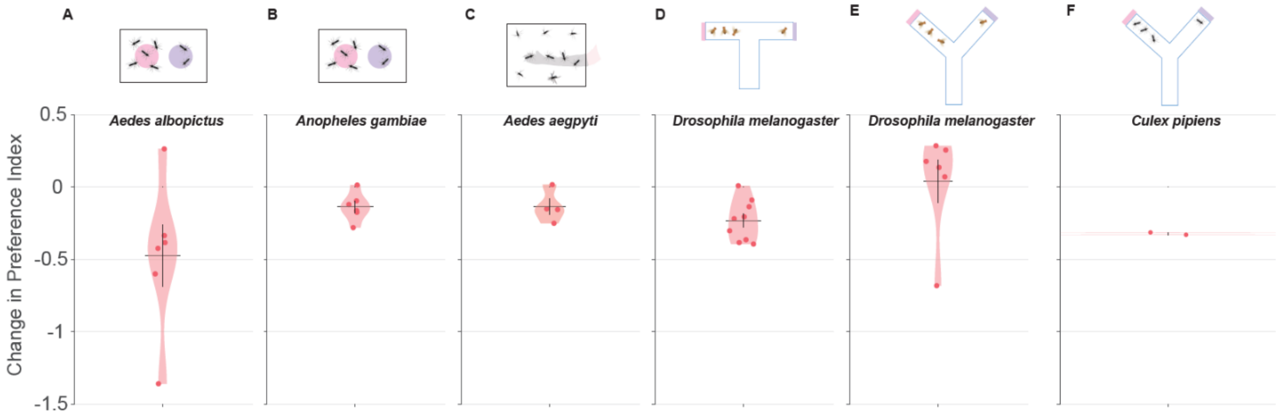


**Fig. S6: Dependence of behavioral preference on odor concentration for different species.**

The change in preference index on increasing the concentration by ten folds for *Aedes albopictus* in dual-port (**A**), *Anopheles gambiae* in dual-port (**B**), *Aedes aegypti* in landing (**C**), *Drosophila melanogaster* in T-maze (**D**), *Drosophila melanogaster* in Y-maze (**E**), and Culex pipiens in Y-maze (**F**) assays.
